# Underreported in-water behaviours of the loggerhead sea turtle: *Getting buried in the sand*

**DOI:** 10.1101/2022.08.26.505133

**Authors:** Kostas Papafitsoros

## Abstract

In this short report, we provide a direct evidence that loggerhead sea turtles *Caretta caretta* are capable of partially burying themselves in the sand by actively moving their front flippers and stirring the sea bottom sediment. In particular, we report the cases of three male loggerhead sea turtles from Zakynthos island, Greece, which, after obtaining a resting position on a sandy spot at the sea bottom, they actively performed digging and stirring movements with their front flippers, resulting to the sand getting raised at the sea column. When the sand settled back, the turtles ended up half-buried and camouflaged. To our current knowledge, this self-burying behaviour has not been described in the literature.

## Introduction

In-water behaviours of sea turtles have been well described in the literature, using a variety of means like direct observations (Booth and Peters, 1972; Schofield *et al*., 2006; Bennett and Keuper-Bennett, 2008), animal-borne cameras (Seminoff *et al*., 2006; Thomson *et al*., 2015), remotely operated vehicles (Smolowitz *et al*., 2015; Dodge *et al*., 2018), static sea floor cameras (Zamzow, 1998) and drones (Schofield *et al*., 2017b). The majority of the studies typically examine standard in-water behaviours like mating (Booth and Peters, 1972; Schofield *et al*., 2006; Schofield *et al*., 2017a), foraging (Smolowitz *et al*., 2015; Wallace *et al*., 2015; Papafitsoros and Schofield, 2016; Schofield *et al*., 2022), avoiding predators (Hounslow *et al*., 2021), cleaning (Zamzow, 1998; Sazima *et al*., 2004; Schofield *et al*., 2017b), and intraspecific interactions (Thomson *et al*., 2015; Gaos *et al*., 2021; Schofield *et al*., 2022). However, since most animals are observed in small time scales, e.g. restricted by the limited battery life of animal-borne cameras, and by several other logistical difficulties, other rare behaviours might remain undetected.

The present note initiates a series of short articles, aiming to describe in-water behaviours of the loggerhead sea turtles (*Caretta caretta*) that are underreported in the literature or not reported at all. Here we report the cases of three male loggerhead sea turtles which, after obtaining a resting position on a sandy spot at the sea bottom, they actively performed digging and stirring movements with their front flippers, resulting to the sand getting raised at the sea column. When the sand settled back, the turtles ended up half-buried and camouflaged. To the author’s current knowledge, this self-burying behaviour has not been reported in the literature.

## Methods

The study site is situated in Laganas Bay, Zakynthos, Greece (37^*◦*^ 43’N, 20^*◦*^ 52’E), Figure 1, which hosts an important breeding site for the Mediterranean loggerhead sea turtles (Casale *et al*., 2018). Long term in-water surveys combined with photoidentification (based on the sea turtles’ unique facial scales) have also shown that Laganas Bay is also a foraging habitat for around 40 resident turtles, mostly males and juveniles (Schofield *et al*., 2020; Papafitsoros *et al*., 2021). The present report is a result of these long term in-water surveys. In summary, surveys involved an observer swimming close to the shore (max 7 metres depth), who, upon encountering a turtle, would take photographs for photo-identification purposes and to record any interesting behaviours. We refer e.g. to (Schofield *et al*., 2020) for more details on the methods and general context.

**Figure 1.**
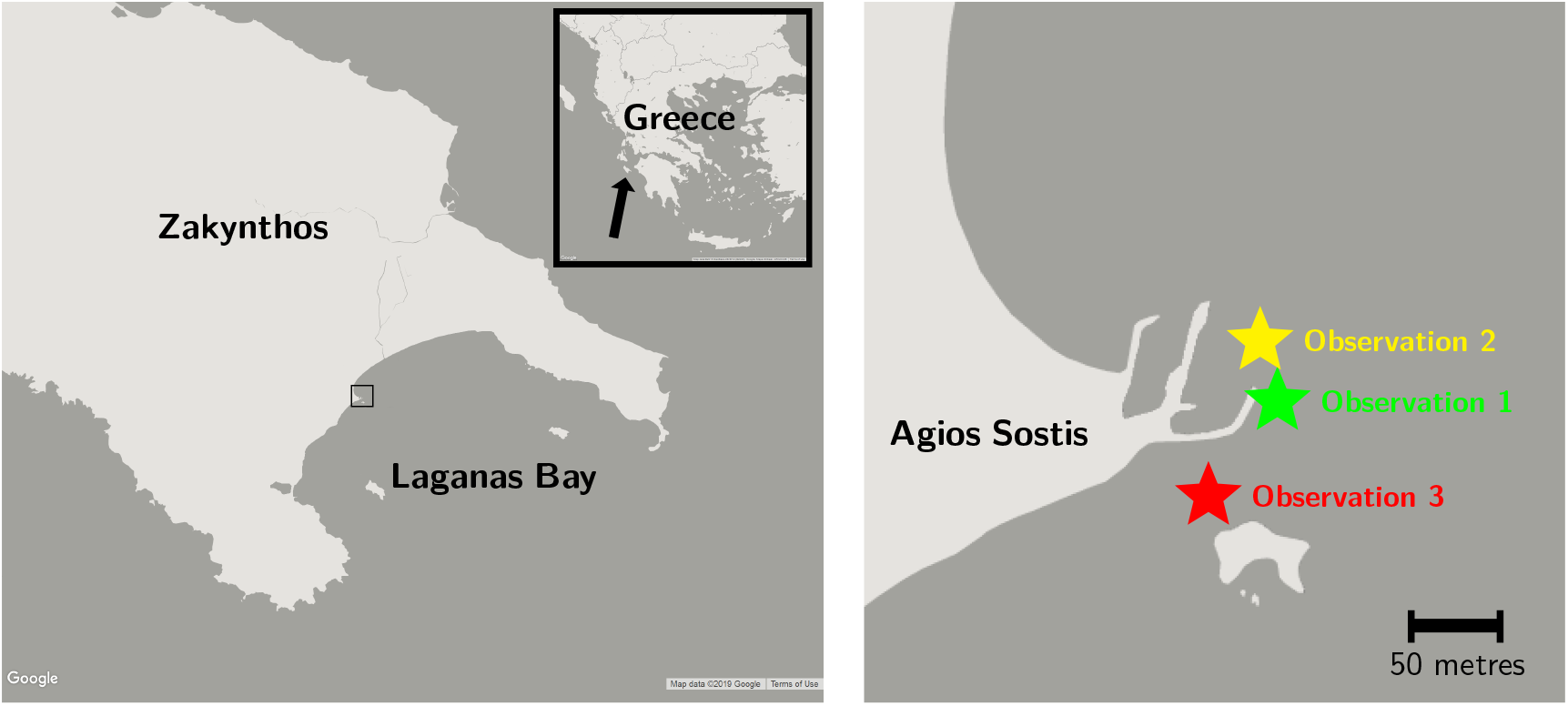
Left: Map of Greece and Laganas Bay, Zakynthos. Right: Map which corresponds to the area enclosed by the small square on the left map, showing the approximate location of the three observations near Agios Sostis area at the northwest part of Laganas Bay.

## Results

All the observations reported next, took place in Agios Sostis area, at the northwest part of Laganas Bay, Zakynthos, see Figure 1 for the approximate locations. All the male turtles involved in Observations 1, 2 and 3, described below, have been long term residents of Laganas Bay, and they were up to a certain degree accustomed to human presence, since they had been regularly observed by tourists (Papafitsoros, 2015; Papafitsoros *et al*., 2021). The first two turtles were adult males (they have been recorded mating), while the third one was a juvenile, based on the length of its tail (Schofield *et al*., 2020).

### Observation 1

This observation took place in 2 October 2016. It began when an adult male loggerhead turtle (ID name “t033”) was approached at 16:44 by a second male that was up to that point being observed. Turtle “t033” had been resting on the sea floor (approximate depth 6 metres) and the presence of the second male triggered an aggressive interaction, with “t033” attacking the second male. The fight lasted approximately 1.5 minutes with the second male fleeing the area. After the fight, “t033” was swimming in the area for about 10 minutes also taking several breaths. At about 16:56, the turtle approached a sandy spot at the sea floor next to a large rock, on which dead seagrass leaves *Posidonia oceanica* were also deposited. The sand was also mixed with oil, something that was inferred from its black colour. This is attributed either to pollution caused by tourist boats that dock to the nearby port or due to some natural oil secretion spots which are common in Laganas Bay. The turtle performed digging and stirring movements with its front flippers for at least 20 seconds, see top part of Figure 2. During that time, the turtle was covered by the cloud-like raised sand and it was essentially not visible. The turtle remained still as the sand settled down and it ended up with its head buried under the seagrass leaves, potentially as it was also slightly moving forward. Sand and debris was deposited on top of its carapace and flippers. The turtle remained still at this position, even upon repeatedly close approaches by the observer (<1 metre), at least until the end of the observation at about 17:00.

**Figure 2.**
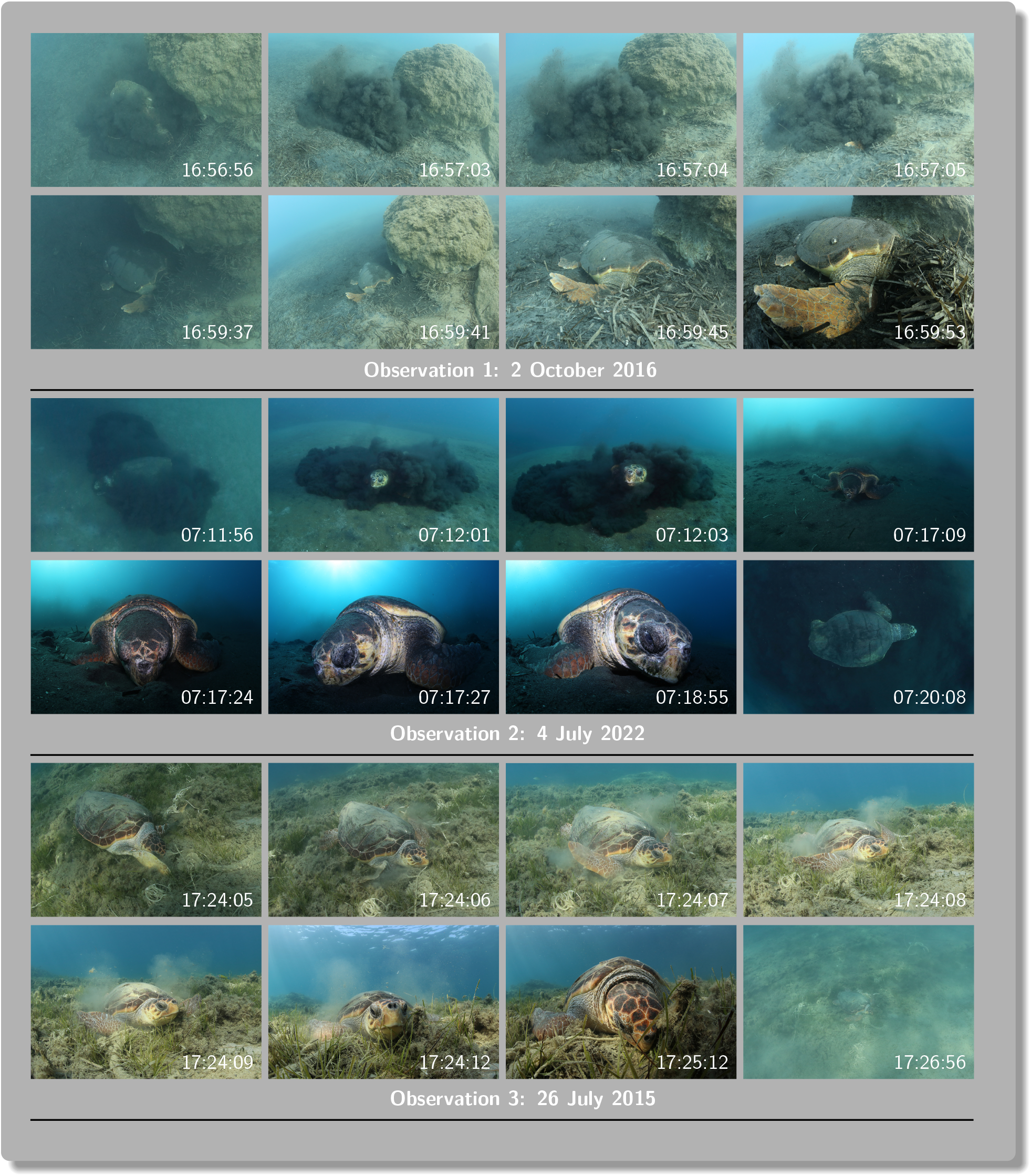
Successive photographs of all the three observations. The times on the bottom left represent the local times that each photograph was taken, as it was internally recorded by the camera.

### Observation 2

This observation took place in 4 July 2022, at an area only a few metres away from the one of Observation 1. The adult male loggerhead “t023” was initially observed swimming in that area at 06:33, trying to forage from the sea floor without success for about 10 minutes. It continued swimming around the area for about 20 minutes. At about 07:11, the turtle approached a spot at the black, again presumably oily, sandy bottom (approximate depth 7 metres) and initiated the same digging and stirring movements also for at least 20 seconds, see middle part of Figure 2. Similarly to Observation 1, the turtle was barely visible during that stage and remained still as the sand settled down. The turtle remained at this position with its carapace, head and flippers covered in sand, at least until the end of the observation at about 07:20.

### Observation 3

This observation took place in 26 July 2015, also at an area close to the ones of the previous observations but at shallower waters (approximate depth 2 metres). Observation of the male “t048” started at 16:36. The turtle was intermittently foraging for sponges until 16:56 where the first part of the observation ended. During that period, it also attacked another foraging male in the area, see Schofield *et al*., 2022 for the nature of these aggressive interactions. The observation resumed at 17:19 and the turtle was swimming around the area until 17:24. At about 17:24 the turtle approached the sea bottom and performed the same front flipper movements as the turtles in Observations 1 and 2. Since the sea bottom consisted of a mix of sand and seagrass, the amount of raised sand was not enough to cover the turtle as much as in the previous cases. Nevertheless the turtle remained still until 17:32 when it came to water surface to breath after which the observation ended. In the meanwhile as in Observation 2, the turtle did not react in an observable manner even after repeatedly close approaches by the observer (<1 metre), but perhaps the last approach at 17:32 triggered the breathing bout.

## Discussion

Getting buried in mud is a well-known behaviour for freshwater turtles during overwintering (Carroll and Ehrenfeld, 1978; Carr, 2018). Sea turtles have also been described to be (half)-buried in sea bottom sediment while being lethargic especially when sea temperature falls below certain thresholds (Felger *et al*., 1976; Carr *et al*., 1980; Ogren and McVea, 1995). In particular, Carr *et al*., 1980 noted that loggerhead sea turtles stuck in the firm sandy clay of Port Canaveral Ship Channel, East Florida, USA, could only be the result of the turtles’ own activity. On the other hand Lamont *et* al., 2021, suggested that sea turtles being buried in sediment is rather a result of them resting on the sea floor for extended periods of times (Hochscheid *et al*., 2005) perhaps facilitated by turbulent water, rather than them actively being buried in the sand. Here we provide a direct evidence that loggerhead sea turtles are indeed capable of getting themselves partially buried in the sand by actively moving their front flippers and stirring the sea bottom sediment. Sea turtles use their flippers for a variety of purposes other than swimming (Fujii *et* al., 2018), and our present observations constitute an additional unreported functioning.

Since sea temperatures in Zakynthos remain well above 20^*◦*^ C from June until October, the observed behaviours reported here cannot be attributed to low temperatures like in the cases mentioned in the references above. Furthermore, the three observed individuals had being in general quite active, e.g. foraging and interacting with other turtles, during the summer periods (author’s personal observations), and also immediately before exhibiting this behaviour. We speculate that self-burying might be a camouflaging behaviour that leads to a decreased chance of getting spotted by other turtles and thus being attacked by them as it is typically done in that site (Papafitsoros and Schofield, 2016; Schofield *et al*., 2022) and/or prevent them from being spotted by predators as well. Uninterrupted resting behaviour can provide an opportunity to conserve energy between foraging events and this self-burying behaviour might just facilitate this.

## References

Bennett, P. & Keuper-Bennett, U. (2008). The book of Honu: Enjoying and learning about Hawaii’s sea turtles. University of Hawaii Press.

Booth, J. & Peters, J.A. (1972). Behavioural studies on the green turtle (Chelonia mydas) in the sea. Animal Behaviour 20(4), 808–812. https://doi.org/10.1016/S0003-3472(72)80155-6.

Carr, A. (2018). Handbook of turtles: the turtles of the United States, Canada, and Baja California. Cornell University Press.

Carr, A., Ogren, L., & McVea, C. (1980). Apparent hibernation by the Atlantic loggerhead turtle Caretta caretta off cape canaveral, Florida. Biological Conservation 19(1), 7–14. https://doi.org/10.1016/0006-3207(80)90011-7.

Carroll, T.E. & Ehrenfeld, D.W. (1978). Intermediate-range homing in the wood turtle, Clemmys insculpta. Copeia (1), 117–126. https://doi.org/10.2307/1443831.

Casale, P., Broderick, A.C., Camiñas, J.A., Cardona, L., Carreras, C., Demetropoulos, A., Fuller, W.J., Godley, B.J., Hochscheid, S., Kaska, Y., Lazar, B., Margaritoulis, D., Panagopoulou, A., Rees, A.F., Tomás, J., & Türkozan, O. (2018). Mediterranean sea turtles: current knowledge and priorities for conservation and research. Endangered species research 36, 229–267. https://doi.org/10.3354/esr00901.

Dodge, K.L., Kukulya, A.L., Burke, E., & Baumgartner, M.F. (2018). TurtleCam: A “smart” autonomous underwater vehicle for investigating behaviors and habitats of sea turtles. Frontiers in Marine Science 5, 90. https://doi.org/10.3389/fmars.2018.00090.

Felger, R.S., Cliffton, K., & Regal, P.J. (1976). Winter dormancy in sea turtles: Independent discovery and exploitation in the Gulf of California by two local cultures. Science 191(4224), 283–285. https://doi.org/10.1126/science.191.4224.283.

Fujii, J.A., McLeish, D., Brooks, A.J., Gaskell, J., & Van Houtan, K.S. (2018). Limb-use by foraging marine turtles, an evolutionary perspective. PeerJ 6, e4565. https://doi.org/10.7717/peerj.4565.

Gaos, A.R., Johnson, C.E., McLeish, D.B., King, C.S., & Senko, J.F. (2021). Interactions among Hawaiian hawksbills suggest prevalence of social behaviors in marine turtles. Chelonian Conservation and Biology 20(2), 167–172. https://doi.org/10.2744/CCB-1481.1.

Hochscheid, S., Bentivegna, F., & Hays, G.C (2005). First records of dive durations for a hibernating sea turtle. Biology Letters 1(1), 82–86. https://doi.org/10.1098/rsbl.2004.0250.

Hounslow, J.L., Jewell, O.J.D., Fossette, S., Whiting, S., Tucker, A.D., Richardson, A., Edwards, D., & Gleiss, A.C. (2021). Animal-borne video from a sea turtle reveals novel anti-predator behaviors. Ecology 102(4), e03251. https://doi.org/10.1002/ecy.3251.

Lamont, M.M, Johnson, D., & Catizone, D.J. (2021). Movements of marine and estuarine turtles during Hurricane Michael. Scientific reports 11, 1577. https://doi.org/10.1038/s41598-021-81234-3.

Ogren, L. & McVea, C. (1995). Apparent hibernation by sea turtles in North American waters. Biology and conservation of sea turtles. Ed. By K.A. Bjorndal. Washington: Smithsonian Institution Press, 127–132.

Papafitsoros, K. (2015). In-water behaviour of the loggerhead sea turtle (Caretta caretta) under the presence of humans (Homo sapiens) in a major Mediterranean nesting site. Proceedings of the 35th annual symposium on sea turtle biology and conservation, Dalaman, Turkey.

Papafitsoros, K., Panagopoulou, A., & Schofield, G. (2021). Social media reveals consistently disproportionate tourism pressure on a threatened marine vertebrate. Animal Conservation 24(4), 568–579. https://doi.org/10.1111/acv.12656.

Papafitsoros, K. & Schofield, G. (2016). Focal photograph surveys: Foraging resident male interactions and female interactions at fish-cleaning stations. Proceedings of the 36th Annual Symposium on Sea Turtle Biology and Conservation, Lima, Peru.

Sazima, Ivan, Grossman, Alice, & Sazima, Cristina (2004). Hawksbill turtles visit moustached barbers: cleaning symbiosis between Eretmochelys imbricata and the shrimp Stenopus hispidus. Biota Neotropica 4, 1–6. https://doi.org/10.1590/S1676-06032004000100011.

Schofield, G.,. Katselidis, K.A., Dimopoulos, P., Pantis, J.D., & Hays, G.C. (2006). Behaviour analysis of the loggerhead sea turtle Caretta caretta from direct in-water observation. Endangered Species Research 2, 71–79. http://dx.doi.org/110.3354/esr002071.

Schofield, G., Katselidis, K.A., Lilley, M.K.S., Reina, R.D., & Hays, G.C. (2017a). Detecting elusive aspects of wildlife ecology using drones: New insights on the mating dynamics and operational sex ratios of sea turtles. Functional Ecology 31(12), 2310–2319. https://doi.org/10.1111/1365-2435.12930.

Schofield, G., Klaassen, M., Papafitsoros, K., Lilley, M., Katselidis, K.A., & Hays, G.C. (2020). Long-term photo-id and satellite tracking reveal sex-biased survival linked to movements in an endangered species. Ecology 101 (7), e03027. https://doi.org/10.1002/ecy.3027.

Schofield, G., Papafitsoros, K., Chapman, C., Shah, A., Westover, L., Dickson, L.C.D., & Katselidis, K.A. (2022). More aggressive sea turtles win fights over foraging resources independent of body size and years of presence. Animal Behaviour 190, 209–219. https://doi.org/10.1016/j.anbehav.2022.05.006.

Schofield, G., Papafitsoros, K., Haughey, R., & Katselidis, K. (2017b). Aerial and underwater surveys reveal temporal variation in cleaning-station use by sea turtles at a temperate breeding area. Marine Ecology Progress Series 575, 153–164. https://doi.org/10.3354/meps12193.

Seminoff, J.A., Jones, T.T., & Marshall, G.J. (2006). Underwater behaviour of green turtles monitored with video-time-depth recorders: what’s missing from dive profiles? Marine Ecology Progress Series 322, 269–280. https://doi.org/10.3354/meps322269.

Smolowitz, R.J., Patel, S.H., Haas, H.L., & Miller, S.A. (2015). Using a remotely operated vehicle (ROV) to observe loggerhead sea turtle (Caretta caretta) behavior on foraging grounds off the mid-Atlantic United States. Journal of Experimental Marine Biology and Ecology 471, 84–91. https://doi.org/10.1016/j.jembe.2015.05.016.

Thomson, J.A., Gulick, A., & Heithaus, M.R. (2015). Intraspecific behavioral dynamics in a green turtle Chelonia mydas foraging aggregation. Marine Ecology Progress Series 532, 243–256. https://doi.org/10.3354/meps11346.

Wallace, B.P., Zolkewitz, M., & James, M.C. (2015). Fine-scale foraging ecology of leatherback turtles. Frontiers in Ecology and Evolution 3. https://doi.org/10.3389/fevo.2015.00015.

Zamzow, J.P. (1998). Cleaning symbioses between Hawaiian reef fishes and green sea turtles, Chelonia mydas. Proceedings of the 18th International Sea Turtle Symposium, Mazatlán, Sinaloa, Mexico.

